# Photometric Decision-Making During the Dawn Choruses of Cicadas

**DOI:** 10.1101/2025.03.10.642322

**Authors:** A. Rakesh Khanna, Raymond E. Goldstein, Adriana I. Pesci, Nir S. Gov

## Abstract

We report the first quantitative study of the onset of dawn choruses of cicadas in several natural habitats. A time-frequency analysis of the acoustical signals is used to define an order parameter for the development of collective singing. The ensemble of recordings reveals that the chorus onset times accurately track the changing sunrise times over the course of many weeks, occurring within civil twilight at a solar elevation of −3.8^°^ ± 0.2^°^. Despite day-to-day variations in the amplitude of fully developed choruses, the order parameter data collapse to a common sigmoidal curve when scaled by those amplitudes and shifted by the onset time, revealing a characteristic rise time of ∼60 s for a chorus to reach saturation amplitude. The results are used to obtain the cumulative distribution function of singing as a function of ground illumination, from which is obtained a generalized susceptibility which exhibits a narrow peak with a half-width of ∼12%. The variance of the order parameter exhibits a similar peak, suggesting that a generalized fluctuation-dissipation theorem holds for this system. A model of decision-making under ramps of a control parameter is developed and can achieve a quantitative match to the data. It suggest that sharpness of the susceptibility peak reflects cooperative decision-making arising from acoustic communication.

---

Among the most familiar collective behaviors in the animal world are the choruses of birds and insects at dawn and dusk. In the case of birds, there is a long history of quantitative observational studies dating back well over a century [1] noting how dawn choruses track the seasonally changing sunrise times [2] and the physiology of birds [3–6], and, more recently, quantifying spatio-temporal aspects of multi-species choruses [7]. Many hypotheses have been advanced to explain *why* birds engage in choruses, with explanations ranging from diurnal variations in individual physiology to ones based on social functions of the choruses [8, 9]. In addition, mechanisms that underlie the observed synchrony of singing have been investigated both for birds and insects [10–15].

Twilight bird choruses are but one example of collective behavior in response to a changing external cue. A spectacular entomological example on longer time scales is the vast swarms of periodical cicadas that emerge after a (prime) number of years spent developing in underground burrows as the soil warms in springtime [16]. Seminal work [17] showed that emergence is associated with the soil temperature —and cicada body temperature—passing through a fairly well-defined threshold value, as evidenced by the staggered emergence over several weeks of subpopulations in markedly different microclimates (sunny south-facing hilltops, shaded valleys).

On closer inspection [18] the problem of decisionmaking under such ramps in an external stimulus is complicated by the significant local variations in soil temperature experienced by underground cicada nymphs even within a particular microclimate due to different burrowing depths and local insolation. It was suggested [18] that communication between underground nymphs, most likely by acoustic signaling, could overcome this environmental noise and lead to larger, more coherent swarms than those that would be produced by individuals responding only to their perceived thermal environment.

Such considerations suggest that these kinds of collective decision-making problems conform to a conceptual picture advanced in the context of human decisionmaking [19, 20]. There, each individual is aware of “public information” that is known to all, perhaps with additive noise, and is coupled to a set of near neighbors who are also in the process of decision-making. An individual’s decision that a threshold in the public information has been crossed is thus affected by its neighbors’ opinions. These ideas naturally lead to theories based on the random-field Ising model in which a uniform external field is slowly changed in time. Examples of processes described by this approach range from coordinated applause at a concert to the mass selling of stocks.

In light of these developments, avian and insectile choruses can serve as paradigms with which to understand decision-making by populations subjected to slowly changing cues. While the vast body of prior work on such choruses mentioned above has almost exclusively focused on defining only the apparent *start* of collective behavior, in reality the choruses grow in amplitude over a finite time scale on the order of minutes. In this paper, in contrast, we focus on the detailed temporal development of the dawn choruses of cicadas with the goal of quantifying how the synchronous singing develops in response to changing light levels. This is done by extracting from the acoustic signals an order parameter for the amplitude of the choruses. We find first that the dawn choruses on clear days commence at a sharply defined value of the pre-dawn solar elevation, which corresponds to a critical ground illumination level *I*, and that, in fact, on cloudy days the onset is delayed. Secondly, the daily chorus amplitudes are found to grow sigmoidally as a function of light intensity and are self-similar. Third, we extract from the chorus amplitudes a generalized susceptibility *χ*(*I*) of singing to light and show that *χ*(*I*) is sharply-peaked around a critical intensity *I*_*c*_. The fluctuations around the average *C* are also found to peak at *I*_*c*_, suggesting the existence of a generalized fluctuation-dissipation theorem at work. Finally, we develop a mathematical model for decision-making that suggests that this sharpness arises from collective effects.

Recordings of the choruses produced by the species *Platypleura capitata* [21] were obtained at two distinct sites near Bangalore, India on multiple days in April and May of 2023, as indicated in Table I. Site I is a shrubland with scattered grasses and Site II is a bamboo forest. Figure 1 displays photographs of the terrain and a typical example of *P. capitata*.

**TABLE I.** Recordings locations and dates.

| Site | Coordinates | Acquisition Dates |
| --- | --- | --- |
| I | 13°9'18.82"N, 77°28'18.04"E | 24 April-1 May, 2023 |
| II | 13°9'28.63"N, 77°29'48.99"E | 15-20 May, 2023 |

**FIG. 1.**
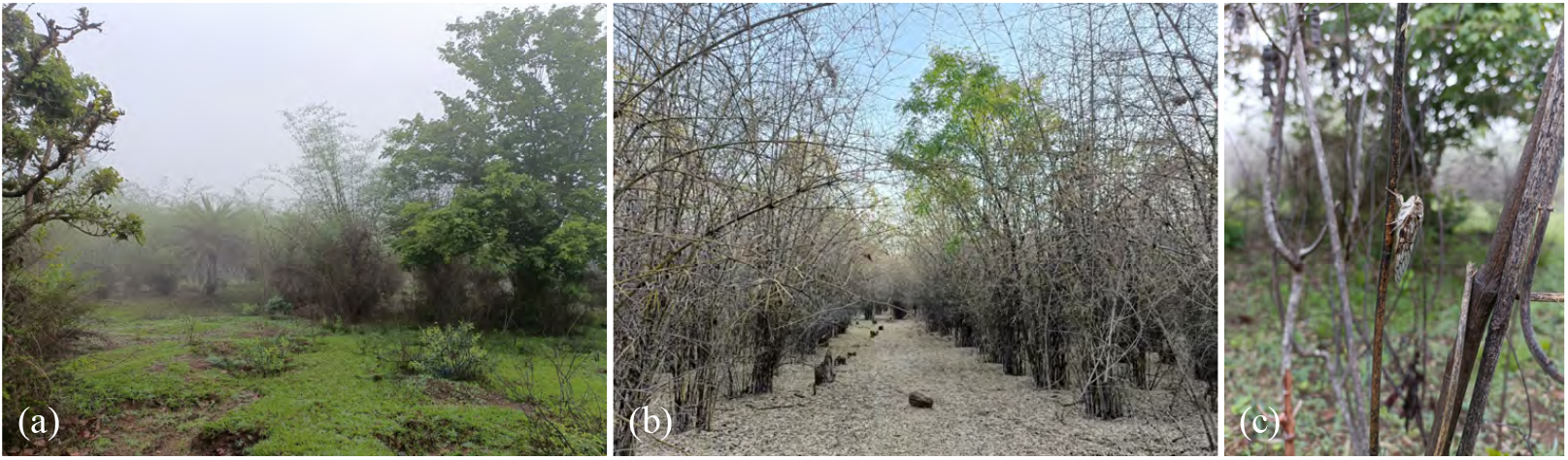
Field observations. (a,b) Sites I and II, displaying range of flora. (c) Exemplar of *P. capitata* on a tree.

Stereo recordings were obtained at a sampling rate of 24 kHz using an off-the-shelf recorder whose Nyquist frequency of 12 kHz is approximately twice the dominant frequencies found in the choruses (see below), assuring that they are accurately captured. The recorder was mounted ∼ 1 m above the ground on a tree branch and remained in a single place for 8 continuous days of recording at each site, broken up into consecutive time-stamped 4 hour recordings that were spliced together for analysis. Signals were analyzed with built-in and bespoke signal processing algorithms in Matlab.

The recorded signals *S*(*t*) are composed of the episodic cicada choruses superimposed on background noise arising from wind, other insects and distant environmental sounds. A spectral analysis of the 8 dawn choruses at location I, shown in Fig. 2, reveals structure at frequencies below ∼ 4 kHz whose amplitude varies considerably from day to day, and a highly reproducible peak above that frequency, with a clear flat maximum extending from a lower frequency of *f*_*l*_ = 4.3 kHz to an upper frequency *f*_*u*_ = 6.3 kHz, the latter being well below the Nyquist frequency. By analyzing segments of the recordings after the end of each dawn chorus we obtained the power spectra in the absence of singing. Figure 2 shows these data as bands of ± one standard deviation around the mean. It is clear from this that the background noise is three orders of magnitude below the main signal. Based on this observation, we chose for all subsequent analysis to consider the filtered signal 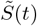 obtained by passing *S*(*t*) through a 20^th^ order Butterworth bandpass filter with lower and upper limits *f*_*l*_ and *f*_*u*_, yielding the truncated average spectrum shown in Fig. 2.

**FIG. 2.**
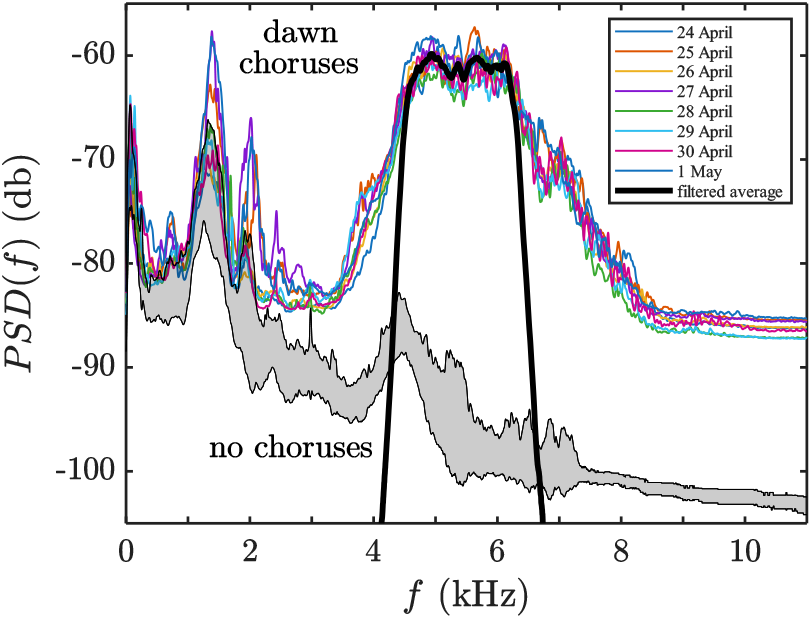
Power spectra of dawn choruses at location I. Colored thin lines are raw power spectral densities of the 8 data sets, while heavy black line is the average of the 8 bandpass-filtered signals. Shaded gray region shows mean ± standard deviation for 8 quiet periods after the choruses.

There is a vast separation of time scales between the sub-millisecond period associated with the dominant chorus pitch and the many seconds over which the amplitude of the chorus develops to a quasi-steady value. Thus, we can obtain a time-dependent measure of the amplitude of a chorus by dividing up the timeline of the signal into intervals of duration *T* (in practice, we use *T* = 5 s) and computing within each interval the power *Q*(*t*) in the filtered signal 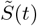 as

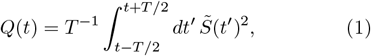

which is a simple implementation of a time-frequency analysis [22]. For the pure tone *S* = *a* sin(2*πf*_0_*t*) with *f*_0_ lying inside the bandpass window, then *Q* = *a*^2^*/*2, and *Q*^1*/*2^ ∝ *a* is a convenient measure of the sound amplitude. We thus define the order parameter as *A*(*t*) = *Q*^1*/*2^(*t*).

Using this procedure, Fig. 3 shows *Q*(*t*) of the daily choruses at site I. Viewed on this coarse time scale, both the dawn and dusk choruses exhibit highly reproducible onset times and more varied termination times. The midday choruses are highly variable in detail from day to day, although they are generally most noticeable during the period 08 : 00 − 16 : 00.

**FIG. 3.**
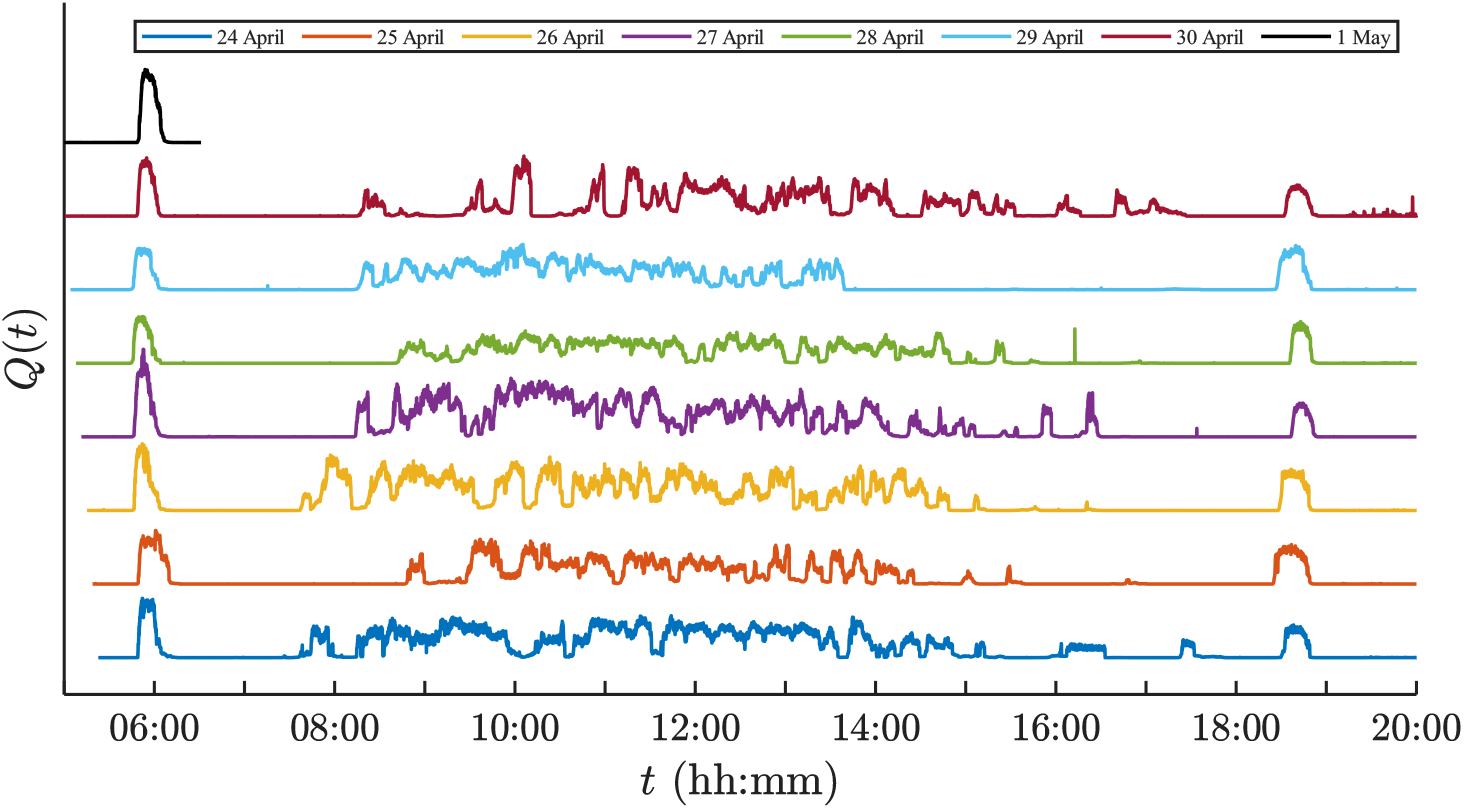
Timeline of daily cicada choruses at location I.

In this paper we focus on the dawn choruses. Figure 4(a) gives a magnified view of the amplitude *A*(*t*) as a function of time at location II, illustrating that the start of each chorus exhibits a similar functional form, albeit with different saturating values *A*_max_(*d*) on each day *d*. The same behavior is found in the dawn choruses at site I. Given these features, it is natural to ask if the data can be collapsed under suitable scalings. For data from both sites I and II, Fig. 4(b) illustrates a test using what we term the *chorus order parameter*

**FIG. 4.**
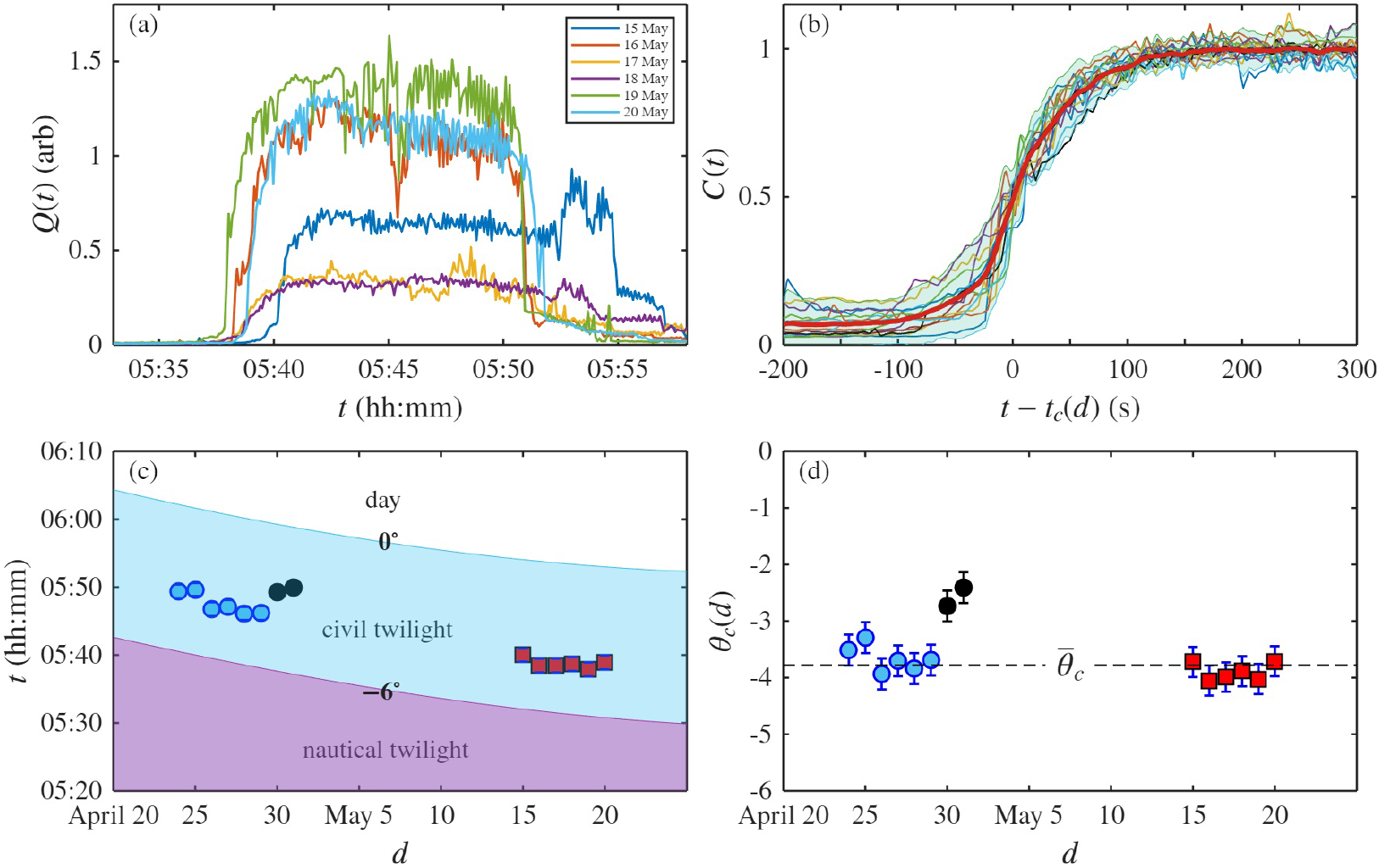
Dawn choruses. (a) Magnified view of dawn chorus power *Q*(*t*) at site II. (b) Chorus order parameter *C*(*t*) near onset, for sites I and II, showing data collapse. Heavy red line is mean, while shaded area represents 95% confidence intervals. (c) Chorus onset times relative to civil twilight (solar elevation of −6^°^) and sunrise (0^°^) for sites I (circles) and II (squares). For site I, light blue circles are for fair weather, black for cloudy days. Smooth boundaries between the different periods are cubic interpolants of public data [23]. (d) Solar elevation at chorus onset versus day. Average 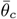 excludes two cloudy days.

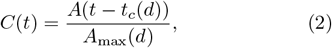

where *t*_*c*_(*d*) is a day-dependent shift termed the *chorus onset time*, defined such that *C*(*t*_*c*_(*d*)) = 1*/*2. We see from the figure that the quality of the data collapse is good, encompassing distinct natural locations several weeks apart. A simple parameterization of the sigmoidal shape is *C*(*t*) = (1 + tanh(*t/τ*))*/*2, from which we obtain the estimate *τ* ∼ 60 s for the characteristic rise time of a chorus. This time is well-resolved given that it is an order of magnitude larger than the sampling window *T*. An interesting feature to which we return below is the markedly larger day-to-day fluctuations around the mean near *t*_*c*_.

We next examine how the chorus onset times *t*_*c*_(*d*) relate to the standard pre-dawn periods that are defined by the ranges of (negative) solar elevation angle *θ*, known as astronomical twilight (−18^°^≤*θ <* −12^°^), nautical twilight (−12^°^≤*θ <* −6^°^) and civil twilight (6^°^≤ *θ <* 0^°^). Data on the beginning and end of each of these periods is readily available [23]. Figure 4(c) shows that *t*_*c*_(*d*) tracks the civil twilight/nautical twilight boundary *t*_*CT*_ (*d*). This is re-expressed in Fig. 4(d) as the solar elevation *θ*_*c*_(*d*) at onset, computed (in degrees) as

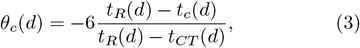

where *t*_*R*_(*d*) is the sunrise time. Examination of historical records on cloud cover [24] shows that the two final days of the data set at site I were significantly cloudy and preceded by heavy rain and thunderstorms, as verified by the audio recordings. This strongly suggests that the delay in the choruses is due to a lower light intensity due to the cloud cover. The records also show several days in which, while the weather was clear in the period leading up to the chorus, rain had occurred many hours earlier, leading to a noticeable increase in humidity and decrease in temperature. Nevertheless, the choruses on those clear days conformed to the general pattern, indicating that temperature and humidity do not play a role in the chorus onset. Excluding the data from site I on those cloudy days, the remaining data cluster very well around the value 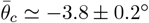.

These results indicate the high precision with which the choruses are associated with a particular solar elevation. This precision can be re-expressed in terms of the solar illumination *I* experienced by the cicadas. Direct measurements of the prevailing illumination with a hand-held light meter showed values of 4 − 6 lux at the beginning of the choruses, and we may appeal to standard astronomical observations [25] to obtain estimates during the entirety of civil twilight. Figure 5(a) shows that the ground illumination *I*(*θ*) varies over ∼ 2.5 orders of magnitude as the sun moves from −6^°^ to 0^°^.

**FIG. 5.**
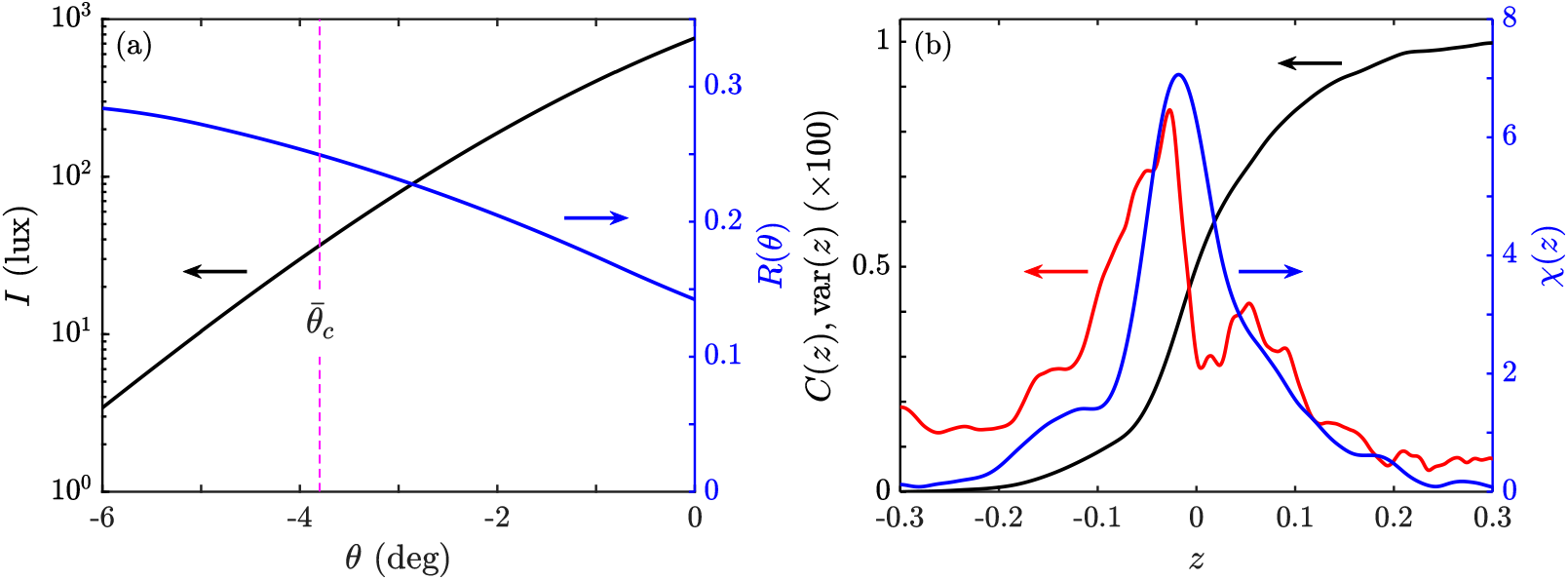
Illumination and decision-making. (a) Ground illumination *I* as a function of solar elevation *θ* (black, left axis) during civil twilight, using standard formulae [25]. Right axis (blue) shows relative changes in illumination from (4) during the period, using *τ* = 60 s. (b) Smoothed values of chorus order parameter *C*(*z*) (black), variance var(*z*) of the order parameter (red) and generalized susceptibility *χ*(*z*) (blue) as functions of the logarithm of normalized ground illumination (5).

In view of the relatively short time scale *τ* ∼ 60 s over which cicada choruses develop, it is natural to estimate the changes in *I* during that period. Earth’s rotational frequency *ω*_*E*_ is 360^°^*/*24 h, or conveniently (1*/*4)^°^ /min. Thus, in the time *τ* ∼ 1 min associated with the onset of the dawn chorus the Earth rotates through an angle *δθ* = *ω*_*E*_*τ* ∼ (1*/*4)^°^. As the rotation is slow compared to the chorus onset time, the relative change *R*(*θ*) in ground illumination during the rise of the chorus can be estimated as

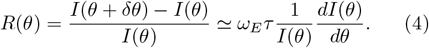

As shown in Fig. 5(a), *R*(*θ*) varies by less than a factor of 2 during civil twilight, with 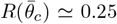. We conclude that under cloudless conditions and in the absence of any obstructing foliage the ground illumination would vary by ∼ 25% during the rise time of the dawn chorus.

A more detailed examination of the photometric response involves viewing the chorus order parameter *C*(*t*) not as a function of time, but rather as a function of the (time-dependent) ground illumination *I*. This is analogous to the way in which the changing time-dependent ground temperature for cicada emergences is the relevant parameter [17, 18]. As it is typical for visual systems to exhibit logarithmic light sensitivity (the Weber-Fechner law [26]), we show in Fig. 5(b) *C* as a function of

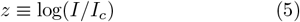

smoothed by a Savitzky-Golay (SG) filter. To make a thermodynamics analogy, we view *C*(*z*) as an order parameter as a function of the variable *z* and hence there is a generalized susceptibility *χ*(*z*) defined as

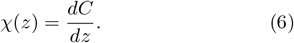

The SG-filtered *χ*(*z*) is shown in blue in Fig. 5(b). The central peak of *χ*(*z*) is reasonably well approximated by a Gaussian, with a standard deviation of 0.06 on a logarithmic scale or 0.14 on a linear scale, reinforcing the conclusion that the singing decision is made with ∼25% precision with respect to the changing illumination.

Also shown in Fig. 5(b) is the SG-filtered variance of the order parameter *C*(*z*). As can be seen in Fig. 4(b), the scale of the standard deviation is ∼ 0.1, so the variance is ∼ 1%. While small, it shows a somewhat noisy, but clear peak near *z* = 0, and in fact closely parallels the susceptibility *χ*(*z*). A variance that is proportional to the susceptibility is evidence of a generalized fluctuation-dissipation theorem [27], where, rather than being derivable from statistical physics, the proportionality constant must be viewed as a distinct property of each given system.

In developing a model for these observations, we begin by noting that the significant variations in the local illumination based on surrounding vegetation levels and daily cicada positions and orientations, is a strong indication that the existence of such a precisely defined transition to the dawn choruses requires a group decisionmaking process. The notion that groups of organisms can make more accurate decisions when acting collectively [28] has been explored in contexts ranging from animal and insect locomotion [29] to discrimination between possible nesting sites of ants [30], bacterial density determination in the process of quorum sensing [31], and insect clocks [32], often using a statistical physics approach based on spin models [33].

A simple model to describe the onset of singing, motivated by the spin description of animal activity and interactions [34], involves assigning to each insect *i* a variable *n*_*i*_, where *n*_*i*_ = 0(1) is the quiet (singing) state. The chorus order parameter is *C* = ⟨*n*⟩. The collection of variables is governed by an energy *E* with an external field 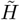 and a coupling 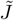,

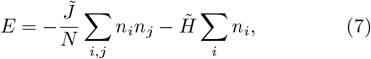

where the light intensity, expressed through the variable *z* in (5), is represented by the external field 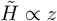. In this way, by itself, a sweep from 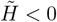 to 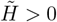 would result in the variables flipping from 0 to 1. Thus, 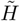 corresponds to the public information, and the infinite-range spin-spin coupling 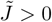 (reflecting acoustic communication as in midge swarms [35]) fosters collective behavior under the assumption that all cicadas behave identically.

In light of the long-ranged coupling between the spins and the fact that *N* ≫ 1, the dynamics of transitions between states can be described through a mean field approximation that takes the form

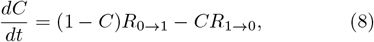

where *R*_0→1_ and *R*_1→0_ are the relevant transition rates from quiet to singing and from singing to quiet. In the Glauber formalism these rates are

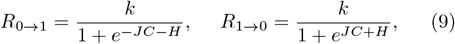

where *k*^−1^ is a decision time of a cicada. Here, *J* and *H* are dimensionless variables corresponding to 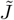 and 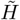 scaled by some effective thermal energy. For *J* = *H* = 0 the two transition rates are both *k/*2. For *J* = 0 the transition to singing is enhanced (and the transition to quiet is diminished) for increasing positive *H*. Similarly, for *J >* 0, the larger the order parameter *C* the more the transition to singing increases; this is the collective effect on decision-making.

When *kτ* ≫ 1 we may use the quasi-static approximation *dC/dt* = 0, giving a self-consistency condition on *C* like that in the theory of ferromagnetism,

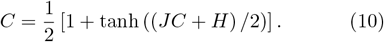

When *J* = 0, (10) yields the sigmoidal curve *C* = (1*/*2)(1 + tanh(*H/*2)); increasing positive *J* produces a more pronounced sigmoid and larger susceptibility. Within this model, the variance of the order parameter can be expressed in a form similar to the rate equation itself (8), as

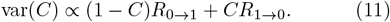

In testing whether the solutions to (10) are consistent with the data in Fig. 5(b), we must recognize that if *H* = *αz* (i.e., proportional rather than strictly equal), then *α* is to be determined along with *J*. Numerical comparison between the field data and the model shows that there is an approximately linear locus in the *α* − *J* plane along which reasonable fits can be obtained, and that along this locus there is a minimum in the total squared deviation between the data and the model (chisquared) at *α* ≃ 15 and *J* ≃ 1.88. A comparison between the data and the optimum of this model is shown in Fig. 6(a), where we see very good agreement with *C*(*z*) and *χ*(*z*). For comparison, the result with *J* = 0 reveals a considerably wider and lower peak in the susceptibility. Figure 6(b) shows that an appropriately scaling of the experimental variance qualitatively matches the theoretical variance. The fact that both *H* and *J* of the optimum model are the same order of magnitude indicates is support for the hypothesis that decision-making in the dawn choruses involves a synergy between external cues and collective effects. In particular, the intermediate value of *J* strikes a balance between on the one hand sharpening the response to changing light levels, giving rise to a rapid chorus development that may be important in such functions as mate attraction [36, 37], and on the other avoiding spontaneous chorus development not strongly correlated with light levels.

**FIG. 6.**
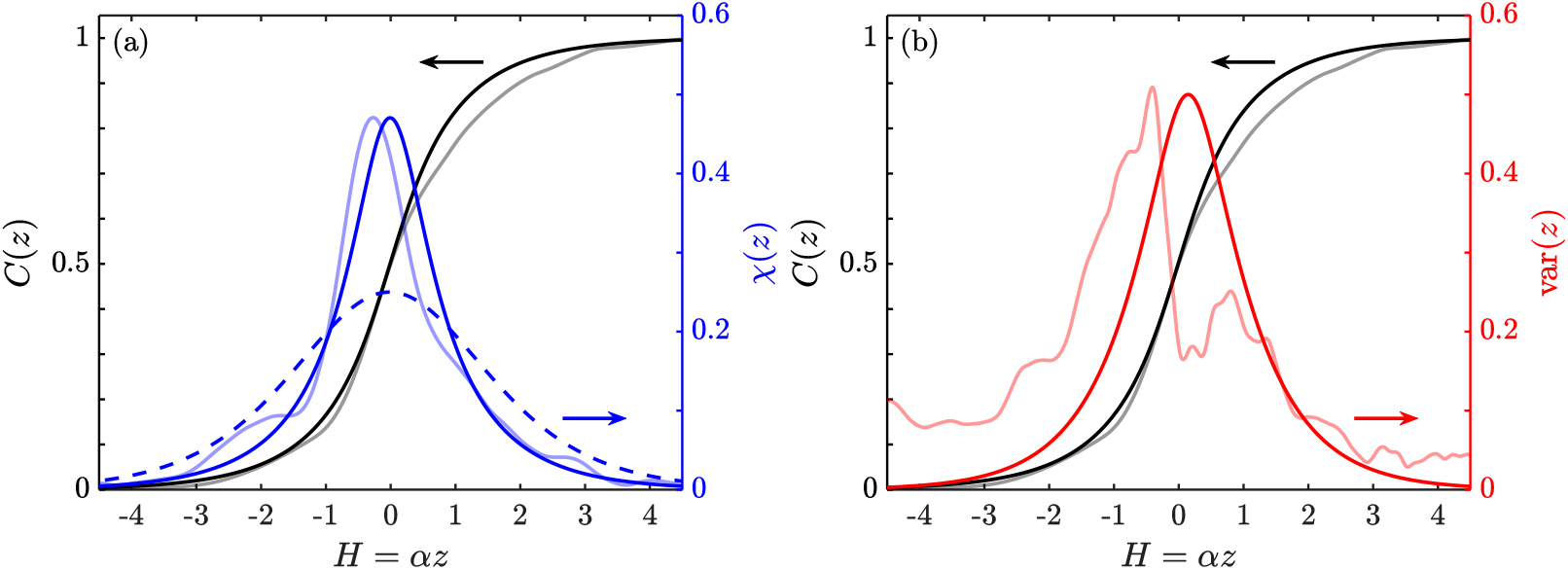
Test of model for decision-making. (a) Chorus order parameter (black) and susceptibility (blue) of optimum model compared with data from Fig. 5(b) (grey, light blue). Dashed blue line indicates susceptibility for *J* = 0. (b) Comparison between model variance (red) and experimental variance (light red), the latter scaled by a factor of 60 for the purposes of comparison.

In this paper we have presented a framework for the analysis of data on collective behavior of decision-making groups insects, using the example of dawn choruses of cicadas. Our work highlights the balance that animals need to reach between individual sensory information (light level) and information shared between individuals (acoustic) in order to optimize their decision-making process. This is a process that occurs generally across collective animal behavior. It is natural to ask whether the methods described here can be applied to a wider group of animals responding to various environmental cues. Further tests of the hypothesis that decisionmaking in the dawn choruses involves communication between equivalent insects could involve longer-term field studies in additional natural habitats, and also interventions to compare the behavior of isolated individuals to that of the group.

## Acknowledgements

We thank Nelson Pesci for inspiring discussions and K. Leptos for helpful comments on the manuscript. This work was supported in part by the Complex Systems Fund at the University of Cambridge (R.E.G. and A.I.P.). N.S.G. is supported by the Lee and William Abramowitz Professorial Chair of Biophysics (Weizmann Institute) with additional support from a Royal Society Wolfson Visiting Fellowship.

The data that support the findings of this article are openly available [38].

## References

[1] C.A. Witchell, The evolution of bird-song: with observations on the influence of heredity and limitation (Adam and Charles Black, London, 1896).

[2] H. Allard, The first morning song of some birds of Washington, D.C.; its relation to light, Am. Nat. 64, 436–469 (1930).

[3] J. Davis, Singing behavior and the gonad cycle of the Rufous-sided Towhee, Condor 60, 308–336 (1958).

[4] A. Leopold and A.E. Eynon, Avian daybreak and evening song in relation to time and light intensity, Condor 63, 269–293 (1961).

[5] R.J. Thomas, T. Székely, I.C. Cuthill, D.G.C. Harper, S.E. Newson, T.D. Frayling and P.D. Wallis, Eye size in birds and the timing of song at dawn, Proc. R. Soc. Lond. B 269, 831–837 (2002).

[6] A. Bruni, D.J. Mennill, and J.R. Foote, Dawn chorus start time variation in a temperate bird community: relationships with seasonality, weather, and ambient light, J. Ornithol. 155, 877–890 (2014).

[7] A. Farina, M. Ceraulo, C. Bobryk, N. Pieretti, E. Quinci and E. Lattanzi, Spatial and temporal variation of bird dawn chorus and successive acoustic morning activity in a Mediterranean landscape, Bioacoustics 24, 269–288 (2015).

[8] C.A. Staicer, D.A. Spector and A.G. Horn The dawn chorus and other diel patterns in acoustic signaling. In: D.E. Kroodsma and E.H. Miller, eds. Ecology and evolution of acoustic communication in birds. Ithaca, NY: Cornell University Press (1996)..

[9] D. Gil, and D. Llusia, The Bird dawn chorus revisited. In Coding Strategies in Vertebrate Acoustic Communication. Animal Signals and Communication 7, T. Aubin and N. Mathevon, eds. (Springer) (2020).

[10] M.D. Greenfield, Synchronous and alternating choruses in insects and aurans: common mechanisms and diverse functions, Amer. Zool. 34, 605–615 (1994).

[11] G. Si-Yuan, J. Yu-Liang, Z. Xiao-Xue, and H. Ji-Ping, A Possible Population-Driven Phase Transition in Cicada Chorus, Comm. Theor. Phys. 51, 1055–1061 (2009).

[12] J. Sueur and T. Aubin, Acoustic communication in the Palaearctic red cicada, Tibicina haematodes: chorus organisation, calling-song structure, and signal recognition, Can. J. Zool. 80, 126–136 (2002).

[13] L.W. Sheppard, B. Mechtley, J.A. Walter, and D.C. Reuman, Self-organizing cicada choruses respond to the local sound and light environment, Ecol. Evol. 10, 4471–4482 (2020).

[14] R. Sarfati, L. Gaudette, J.M. Cicero, and O. Peleg, Statistical analysis reveals the onset of synchrony in sparse swarms of Photinus knulli fireflies, J. R. Soc. Interface 19, 20220007 (2022).

[15] M. McCrea, B. Ermentrout, and J.E. Rubin, A model for the collective synchronization of flashing in photinus carolinus J. Roy. Soc. Interface 19, 20220439 (2022).

[16] C. Simon, J.R. Cooley, R. Karban, and T. Sota, Advances in the Evolution and Ecology of 13- and 17-Year Periodical Cicadas, Annu. Rev. Entomol. 67, 457–482 (2022).

[17] J.E. Heath, Thermal Synchronization of Emergence in Periodical “17-year” Cicadas (Homoptera, Cicadidae, Magicicada), The Am. Mid. Nat. 80, 440–448 (1968).

[18] R.E. Goldstein, R.L. Jack, and A.I. Pesci, How do cicadas emerge together? Thermophysical aspects of their collective decision-making, Phys. Rev. E 109, L022401 (2024).

[19] Q. Michard and J.-P. Bouchaud, Theory of collective opinion shifts: from smooth trends to abrupt swings, Eur. Phys. J. B 47, 151–159 (2005).

[20] J.-P. Bouchaud, Crises and Collective Socio-Economic Phenomena: Simple Models and Challenges, J. Stat. Phys. 151, 567–606 (2013).

[21] https://www.gbif.org/species/8406013

[22] L. Cohen, Time-frequency distributions - a review, Prov. IEEE 77, 941–981 (1989).

[23] The data were obtained from https://www.timeanddate.com/sun/india/bengaluru?month=4&year=2023.

[24] The data were obtained from https://weatherspark.com/h/m/108998/2023/4/Historical-Weather-in-April-2023-in-Bengaluru-Karnataka-India#Figures-CloudCover.

[25] Explanatory Supplement to the Astronomical Almanac, K. Seidelmann, ed., (University Science Books, Mill Valley, CA, 1992), §9.34.

[26] K.O. Johnson, S.S. Hsiao and T. Yoshioka, Neural coding and the basic law of psychophysics, The Neurosci. 8, 111–121 (2002).

[27] K. Sato, Y. Ito, T. Yomo, and K. Kaneko, On the relation between fluctuation and response in biological systems, Proc. Natl. Acad. Sci. (USA) 100, 14086–14090 (2003).

[28] H. Rauhut and J. Lorenz, The wisdom of crowds in one mind: How individuals can simulate the knowledge of diverse societies to reach better decisions, J. Math. Psych. 55, 191–197 (2011).

[29] I.D. Couzin, Collective Cognition in Animal Groups, Trends Cog. Sci. 104, 36–43 (2008).

[30] T. Sasaki, B. Granovskiy, R.P. Mann, D.J.T. Sumpter, and S.C. Pratt, Ant colonies outperform individuals when a sensory discrimination task is difficult but not when it is easy, Proc. Natl. Acad. Sci. USA 110, 13769–13773 (2013).

[31] S. Moreno-Gímez, M.E. Hochberg and G.S. van Doorn, Quorum sensing as a mechanism to harness the wisdom of the crowds, Nat. Comm. 14, 3415 (2023).

[32] G.B.S. Rivas, L.G.S. da R. Bauzer and A.C.A. Meireles-Filho, “The environment is everything that isn’t me”: Molecular mechanisms and evolutionary dynamics of insect clocks in variable surroundings, Front. Physiol.. 6, 400 (2016).

[33] A.T. Hartnett, E. Schertzer, S.A. Levin, and I.D. Couzin, Heterogeneous preference and local nonlinearity in consensus decision making, Phys. Rev. Lett. 116, 038701 (2016).

[34] I. Pinkoviezky, I.D. Couzin, and N.S. Gov, Collective conflict resolution in groups on the move, Phys. Rev. E 97, 032304 (2018).

[35] E. Gorbonos, R. Ianconescu, J.G. Puckett, R. Ni, N.T. Ouellette and N.S. Gov, Long-range acoustic interactions in insect swarms: an adaptive gravity model, New J. Phys. 18, 073042 (2016).

[36] M. Hartbauer and H. Römer, Rhythm generation and rhythm perception in insects: The evolution of synchronous choruses, Front. Neurosci. 10, 223 (2016).

[37] G. Beauchamp, Chorus organization in a neotropical forest cicada, Biology 13, 913 (2024).

[38] R. Khanna A., R.E. Goldstein, A.I. Pesci, and N. Gov, Data for “Photometric decision-making during the dawn choruses of cicadas”. Zenodo. 10.5281/zenodo.14960413 (2025).

